# The macaque brain ONPRC18 template with combined gray and white matter labelmap for multimodal neuroimaging studies of nonhuman primates

**DOI:** 10.1101/2020.10.06.323063

**Authors:** Alison R Weiss, Zheng Liu, Xiaojie Wang, William A Liguore, Christopher D. Kroenke, Jodi L. McBride

**Affiliations:** Division of Neuroscience, Oregon National Primate Research Center, Beaverton, OR, USA, 97006; Advanced Imaging Research Center, Oregon Health and Science University, Portland, OR, USA, 97239; Departments of Behavioral Neuroscience, Portland OR, USA, 97239; Neurology, Oregon Health and Science University, Portland OR, USA, 97239

**Author notes:** **Corresponding author:** Jodi L. McBride, PhD, Associate Professor, Division of Neuroscience, Oregon National Primate Research Center, Department of Behavioral Neuroscience, Oregon Health and Science University, 505 NW 185^th^ Avenue, Beaverton, OR 97006, USA.

**Keywords:** macaque, brain atlas, DTI, MRI, cortico-basal ganglia networks

## Abstract

Macaques are the most common nonhuman primate (NHP) species used in neuroscience research. With the advancement of many neuroimaging techniques, new studies are beginning to apply multiple types of in vivo magnetic resonance imaging (MRI), such as structural imaging (sMRI) with T1 and T2 weighted contrasts alongside diffusion weighed (DW) imaging. In studies involving rhesus macaques, this approach can be used to better understand micro-structural changes that occur during development, in various disease states or with normative aging. However, many of the available rhesus brain atlases have been designed for only one imaging modality, making it difficult to consistently define the same brain regions across multiple imaging modalities in the same subject. To address this, we created a brain atlas from 18 adult rhesus macaques that includes co-registered templates constructed from images frequently used to characterize macroscopic brain structure (T2/SPACE and T1/MP-RAGE), and a diffusion tensor imaging (DTI) template. The DTI template was up-sampled from 1 mm isotropic resolution to resolution match to the T1 and T2-weighted images (0.5 mm isotropic), and the parameter map was derived for fractional anisotropy (FA). The labelmap volumes delineate 57 gray matter regions of interest (ROIs; 36 cortical regions and 21 subcortical structures), as well as 74 white matter tracts. Importantly, the labelmap overlays both the structural and diffusion templates, enabling the same regions to be consistently identified across imaging modalities. A specialized condensed version of the labelmap ROIs are also included to further extend the usefulness of this tool for imaging data with lower spatial resolution, such as functional MRI (fMRI) or positron emission tomography (PET).

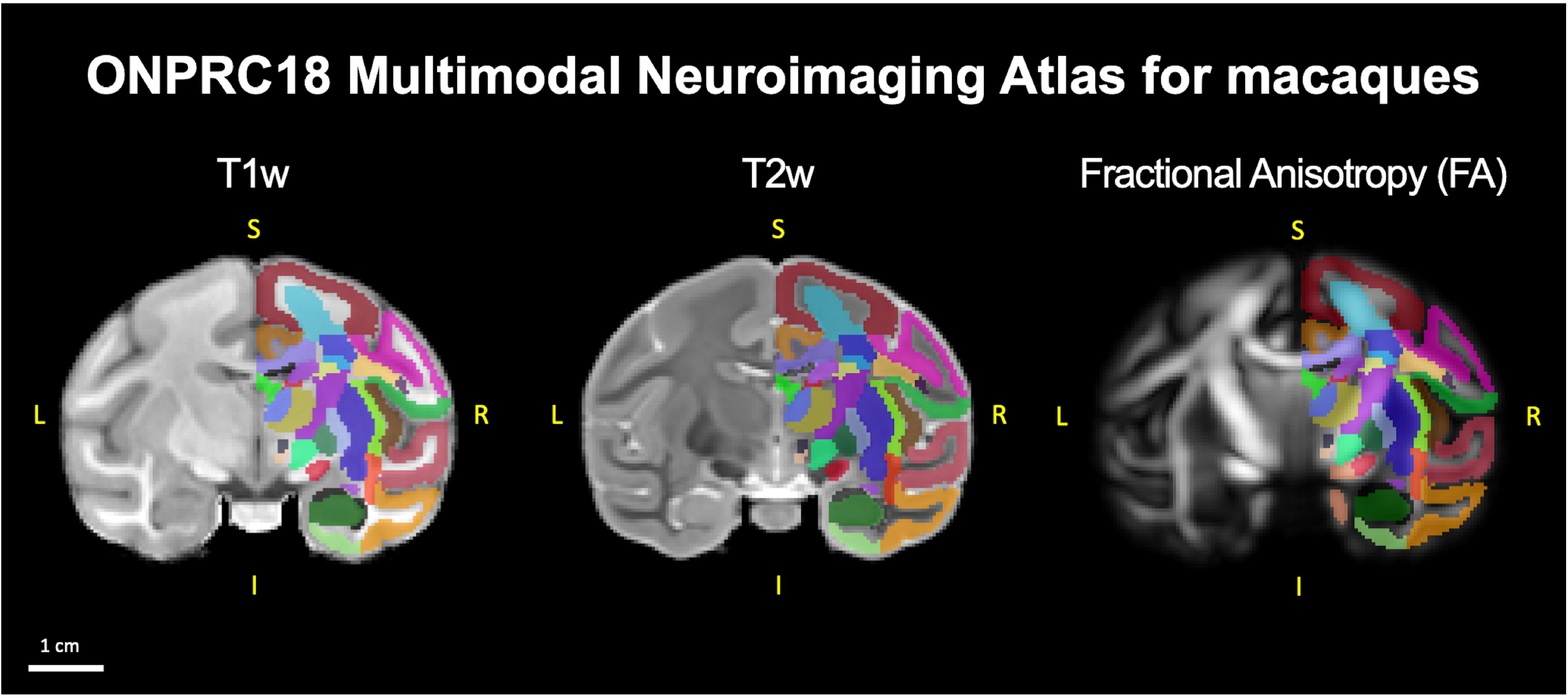
Graphical Abstract.

## 1 INTRODUCTION

Nonhuman primates (NHPs) are valuable for translational neuroscience due to their relatively large and complex brain structure, their diverse behavioral repertoire, and their close phylogenic relationship with humans. Many neuroimaging studies with macaques require a species-specific magnetic resonance imaging (MRI) atlas to accurately delineate the anatomical boundaries of brain regions. Recent advances in neuroimaging techniques have spurred the generation of several NHP-specific MRI atlases and labelmaps, for example a T1-weighted MRI template by Rohlfing et al. (2012), a T2-weighted MRI template by Calabrese et al. (2015), and diffusion tensor imaging (DTI) templates by Zakszewski et al. (2014) and Adluru et al. (2012). Historically, atlases that have been designed for only one imaging modality represent a challenge for studies employing multiple neuroimaging modalities in the same study population. For such multi-modal neuroimaging studies, different templates can be applied separately to each scan modality. However, a limitation to this approach is that the corresponding label maps from each template may have slight-to-significant differences in boundary definitions, making it difficult to consistently define the same brain regions across multiple imaging modalities in the same subject.

To address this gap, we created a series of co-registered templates, including T1- and T2-weighted MRI as well as DTI. Regions of Interest (ROI) were defined to facilitate the investigation of cortico- and thalamo-basal ganglia circuitry and were informed by the boundaries described by Saleem & Logothetis (2007). This atlas builds on previous work from our group (Rohlfing et al., 2012) using non-linear spatial normalization techniques (Avants et al., 2008; Yeo et al., 2008) to generate rhesus brain templates, and integrates high-resolution images from multiple image contrasts.

We have also made practical improvements in the labelmap to facilitate multi-modal neuroimaging studies. The INIA19 atlas (Rohlfing et al., 2012) applied NeuroMaps labels that were originally defined on a histological dataset (Braininfo, 1991-present). However, in practice, we have found that both scan quality and individual variability in neuroanatomy impact the reliability and accuracy of registration-based segmentation approaches using highly-detailed and complex labelmaps with small ROIs. Here, we created a labelmap with fewer, more simplified ROIs that are better matched to the image resolution of in vivo MRI and more tolerant of individual variation. To facilitate use of these templates for multimodal neuroimaging studies, the T1-weighted, T2-weighted, and diffusion-based images are registered to a common reference frame and labelmap. Moreover, to extend the usefulness of this tool for imaging data with lower spatial resolution, such as positron emission tomography (PET) or functional MRI (fMRI), a specialized condensed version of the labelmap ROIs was also generated.

## 2 MATERIALSAND METHODS

### 2.1 Subjects

Eighteen male (n=6) and female (n=12) adult rhesus macaques (ages 5-12) contributed to this study **(Table 1**). At the time the scans were collected, the animals were pair housed in standard indoor caging, maintained on a 7am/7pm light/dark cycle, given ad libitum access to water, and provided with monkey chow rations and fresh produce daily. The Institutional Animal Care and Use Committee (IACUC) approved all procedures used in this study, and all of the guidelines specified in the National Institutes of Health Guide for the Care and Use of Laboratory Animals were strictly followed.

**Table 1:**
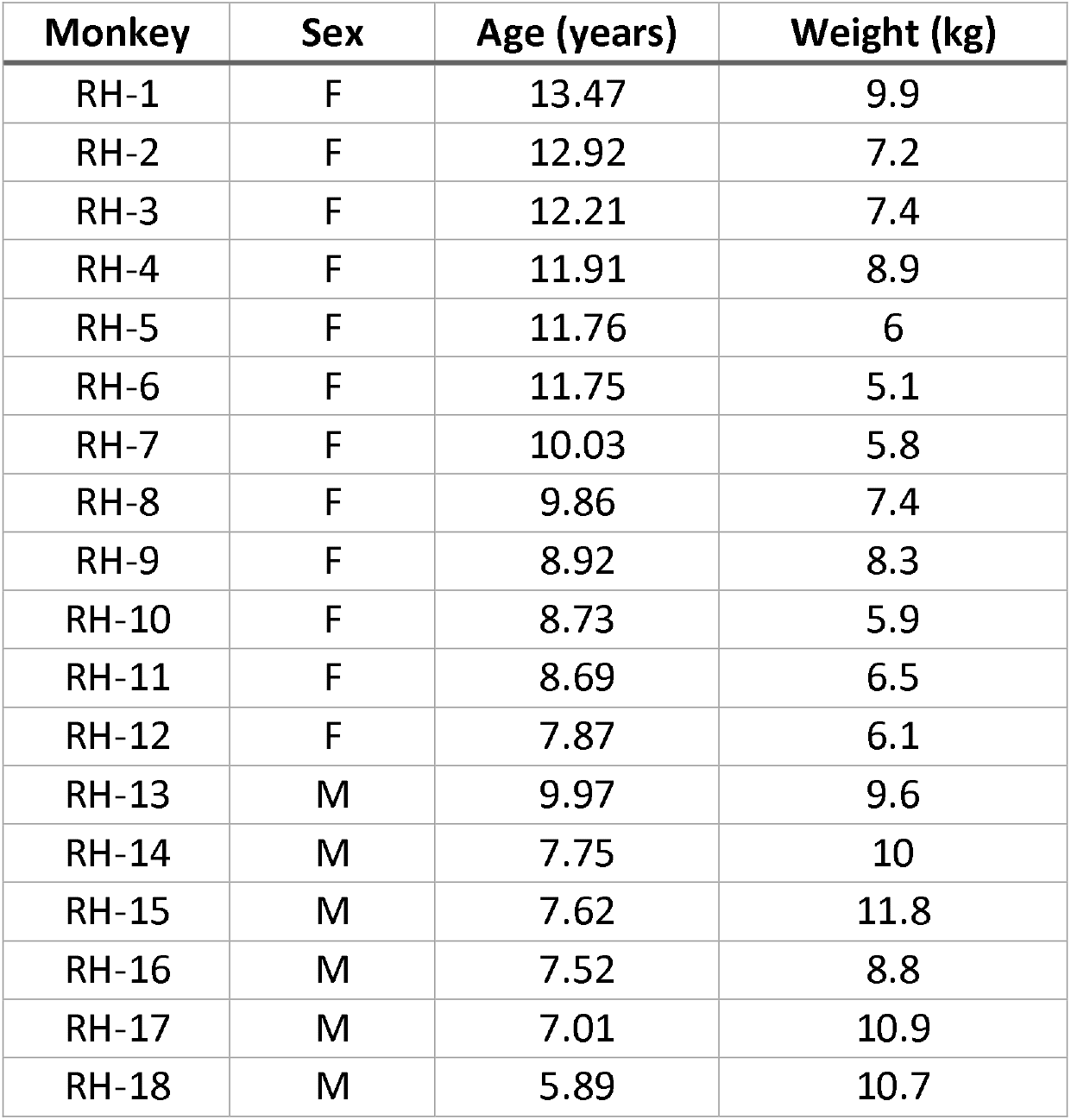
Demographic information for study animals. Scans from 18 individual rhesus macaques contributed to these templates. At the time of the scans, the animals were 6-13 years old, and consisted of both males (n=6) and females (n=12). There were slight age differences between the sexes, with females a bit older than males.

**Table 2:**
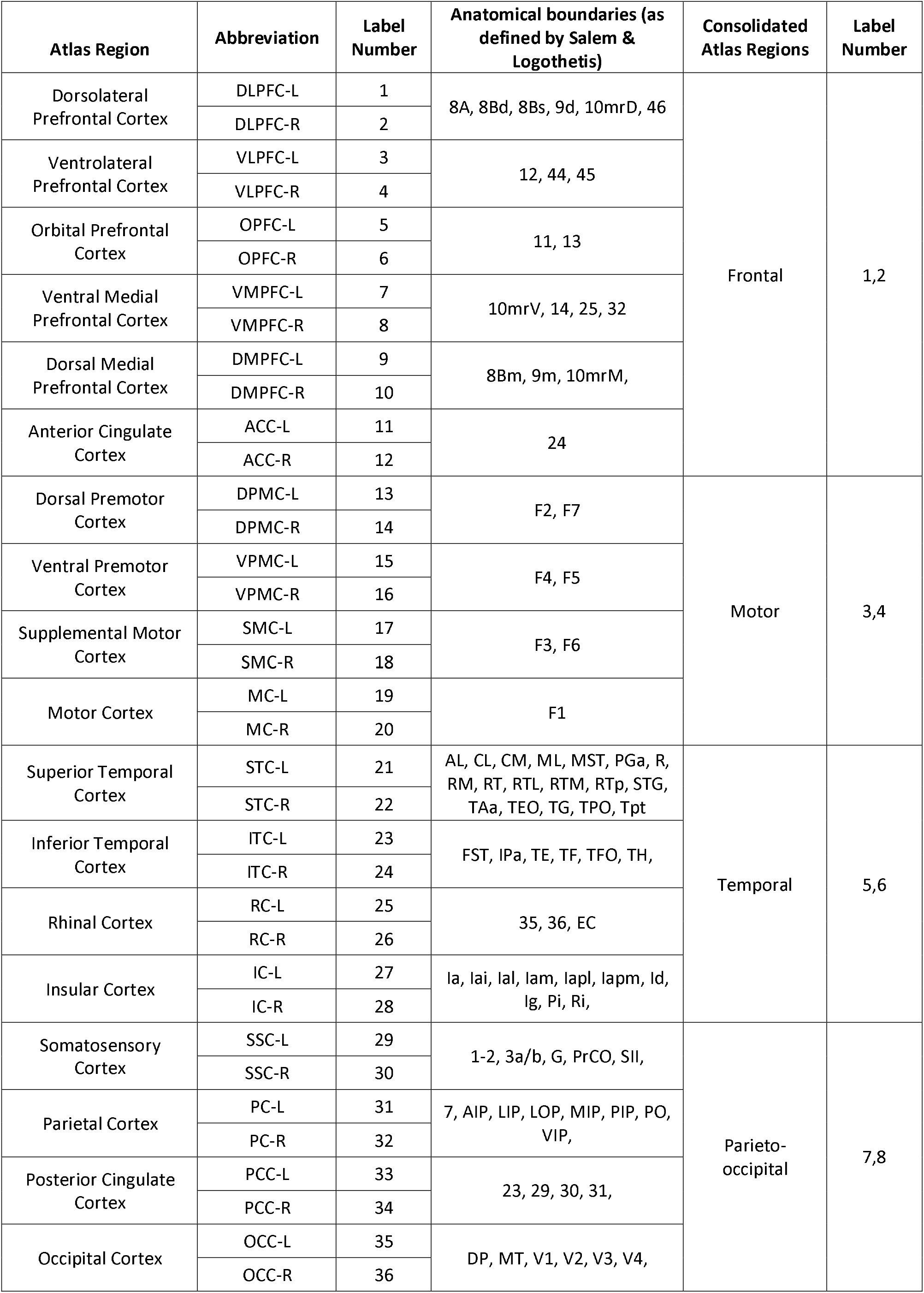
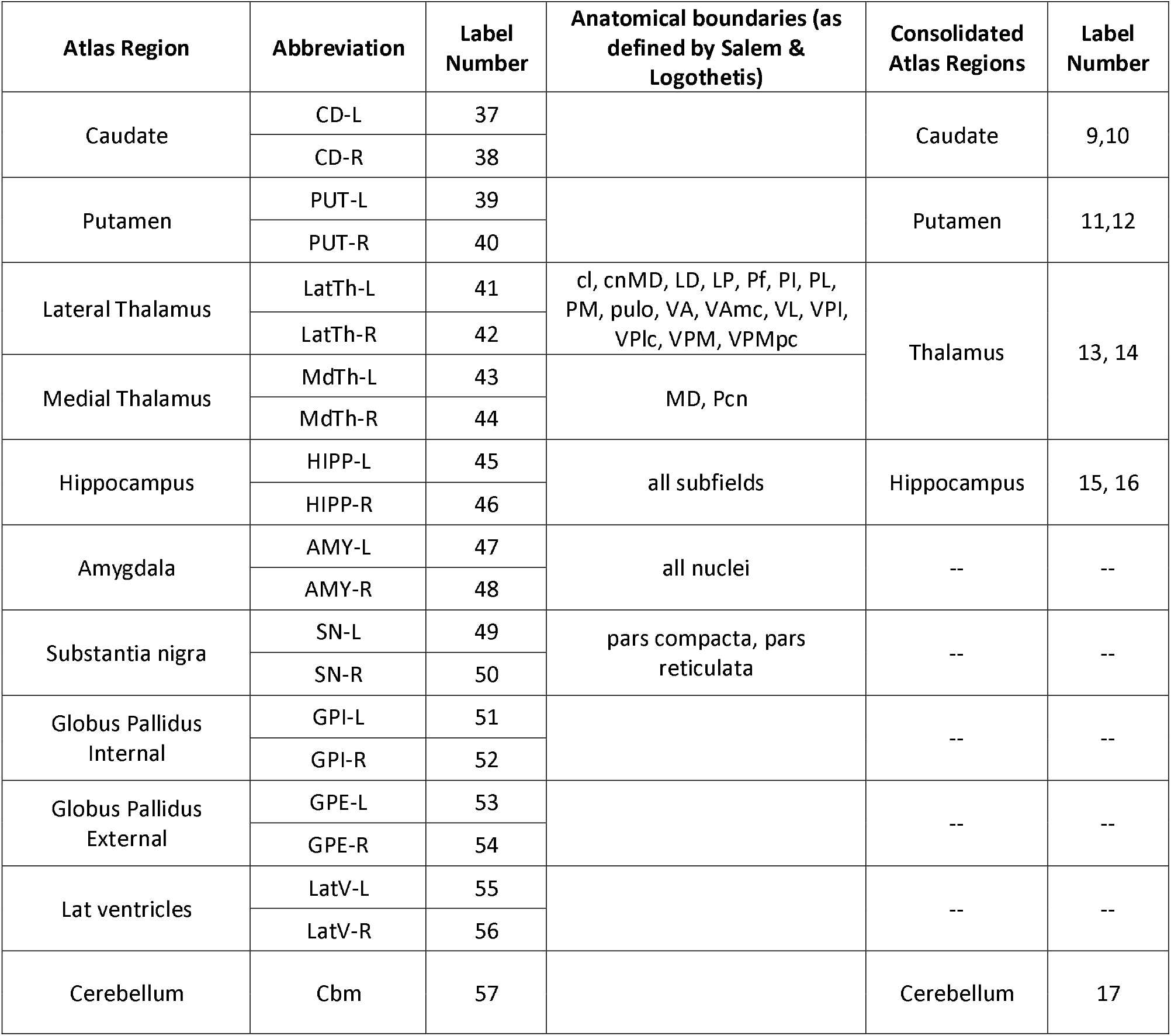
Description of ROI anatomical boundaries in labelmaps. All regions were drawn by hand on the T2 template using the 3D segmentation software ITK-SNAP (Yushkevich et al., 2006). Cortical and subcortical ROIs were defined on coronal sections according to the atlas by Saleem & Logothetis (2007), and verified in the axial and sagittal planes. Regions from the labelmap were subsequently categorized and combined to create a consolidated labelmap that defines 8 bilateral cortical ROIs and 9 subcortical structures. Abbreviations: *1-2*, somatosensory areas 1 and 2; *3a/b*, somatosensory areas 3a and 3b; 7, area 7 (all subdivisions); *8A*, area 8A; *8Bd*, area 8B, dorsal subdivision; *8Bm*, area 8B, medial subdivision; *8Bs*, area 8B in the arcuate sulcus; *9d*, area 9, dorsal subdivision; *9m*, area 9, medial subdivision; *10mrD*, area 10mr dorsal subdivision; *10mrM*, area 10mr medial subdivision; *10mrV*, area 10mr ventral subdivision; *11l*, area 11l; *11m*, area 11m; *12l*, area 12l; *12m*, area 12m; *12*, area 12 (all subdivisions); *13*, area 13 (all subdivisions); *14*, area 14 (all subdivisions); *23*, area 23 (all subdivisions); *24*, area 24 (all subdivisions); 25, area 25; *29*, area 29 (retrosplenial cortex); *30*, area 30 (retrosplenial cortex); *31*, area 31; *32*, area 32; *35*, area 35; *36*, area 36 (all subdivisions); *44*, area 44; 45, area 45; *46*, area 46 (all subdivisions); *AIP*, anterior intraparietal area; *AL*, anterior lateral, belt region of the auditory cortex; *CL*, caudal lateral, belt region of the auditory cortex; *cl*, central lateral nucleus; *CM*, caudomedial, belt region of the auditory cortex; *cnMD*, centromedian nucleus; *DP*, dorsal prelunate area; *EC*, entorhinal cortex (all subdivisions); *F1*, agranular frontal area F1; *F2*, agranular frontal area F2; *F3*, agranular frontal area F3; *F4*, agranular frontal area F4; *F5*, agranular frontal area F5; *F6*, agranular frontal area F6; *F7*, agranular frontal area F7; *FST*, floor of superior temporal area; *G*, gustatory cortex; *1a*, agranular insula; *lal*, intermediate agranular insula area; *Ial*, lateral agranular insula area; *Iam*, medial agranular insula area; *Iapl*, posterolateral agranular insula area; *Iapm*, posteromedial agranular insula area; *Id*, dysgranular insula; *Ig*, granular insula; *IPa*, area IPa (sts fundus); *LD*, lateral dorsal nucleus; *LIP*, lateral intraparietal area (all subdivisions); *LOP*, lateral occipital parietal area; *LP*, lateral posterior nucleus; *MD*, medial dorsal nucleus (all subdivisions); *MIP*, medial intraparietal area; *ML*, middle lateral, belt region of the auditory cortex; *MST*, medial superior temporal area; *MT*, middle temporal area; *Pcn*, paracentral nucleus; *Pf*, parafascicular nucleus; *PGa*, area PGa; *PI*, inferior pulvinar; *Pi*, parainsular area; *PIP*, posterior intraparietal area; *PL*, lateral pulvinar; *PM*, medial pulvinar; *PO*, parieto-occipital area; *PrCO*, precentral opercular area; *pulo*, pulvinar oralis nucleus; *R*, rostral, core region of the auditory cortex; *Ri*, retroinsula; *RM*, rostromedial, belt region of the auditory cortex; *RT*, rostrotemporal, core region of the auditory cortex; *RTL*, lateral rostrotemporal, belt region of the auditory cortex; *RTM*, medial rostrotemporal, belt region of the auditory cortex; *RTp*, rostrotemporal (polar); *SII*, secondary somatosensory area (S2); *STG*, superior temporal gyrus; *TAa*, area TAa (sts dorsal bank); *TE*, area TE (all subdivisions); *TEO*, area TEO; *TF*, area TF of the parahippocampal cortex; *TFO*, area TFO of the parahippocampal cortex, *TG*, area TG, temporal pole (all subdivisions); *TH*, area TH of the parahippocampal cortex; *TPO*, area TPO (sts dorsal bank); *Tpt*, temporo-parietal area; *V1*, visual area 1 (primary visual cortex); *V2*, visual area 2; *V3*, visual area 3 (all subdivisions); *V4*, visual area 4 (all subdivisions); *VA*, ventral anterior nucleus; *VAmc*, ventral anterior nucleus, magnocellular division; *VIP*, ventral intraparietal area; *VL*, ventral lateral nucleus (all subdivisions); *VPI*, ventral posterior inferior nucleus; *VPLc*, ventral posterior lateral caudal nucleus; *VPM*, ventral posterior medial nucleus; *VPMpc*, ventral posterior medial nucleus, parvicellular division. Abbreviations from the atlas of Saleem and Logothetis (2007).

**Table 3:**
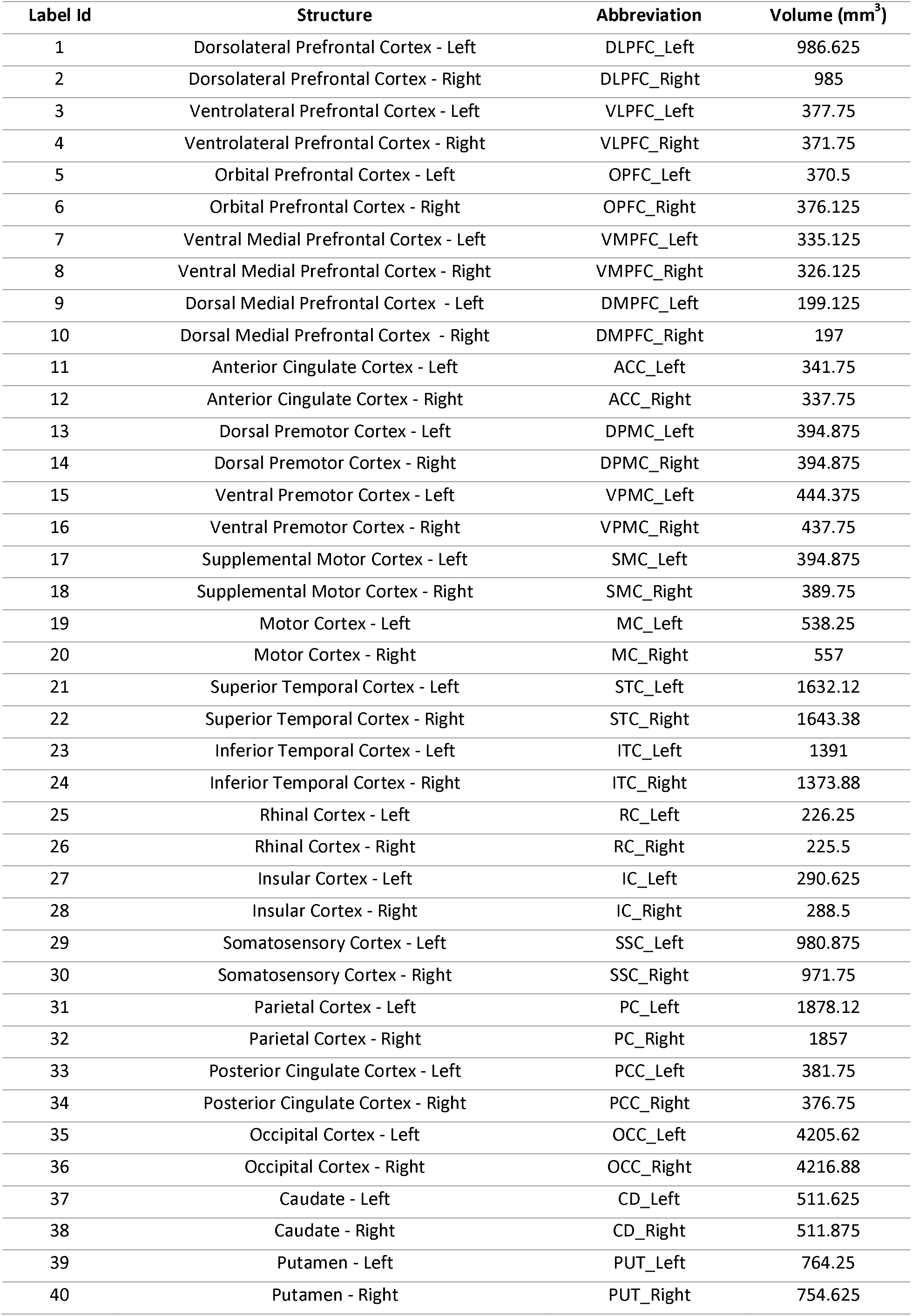
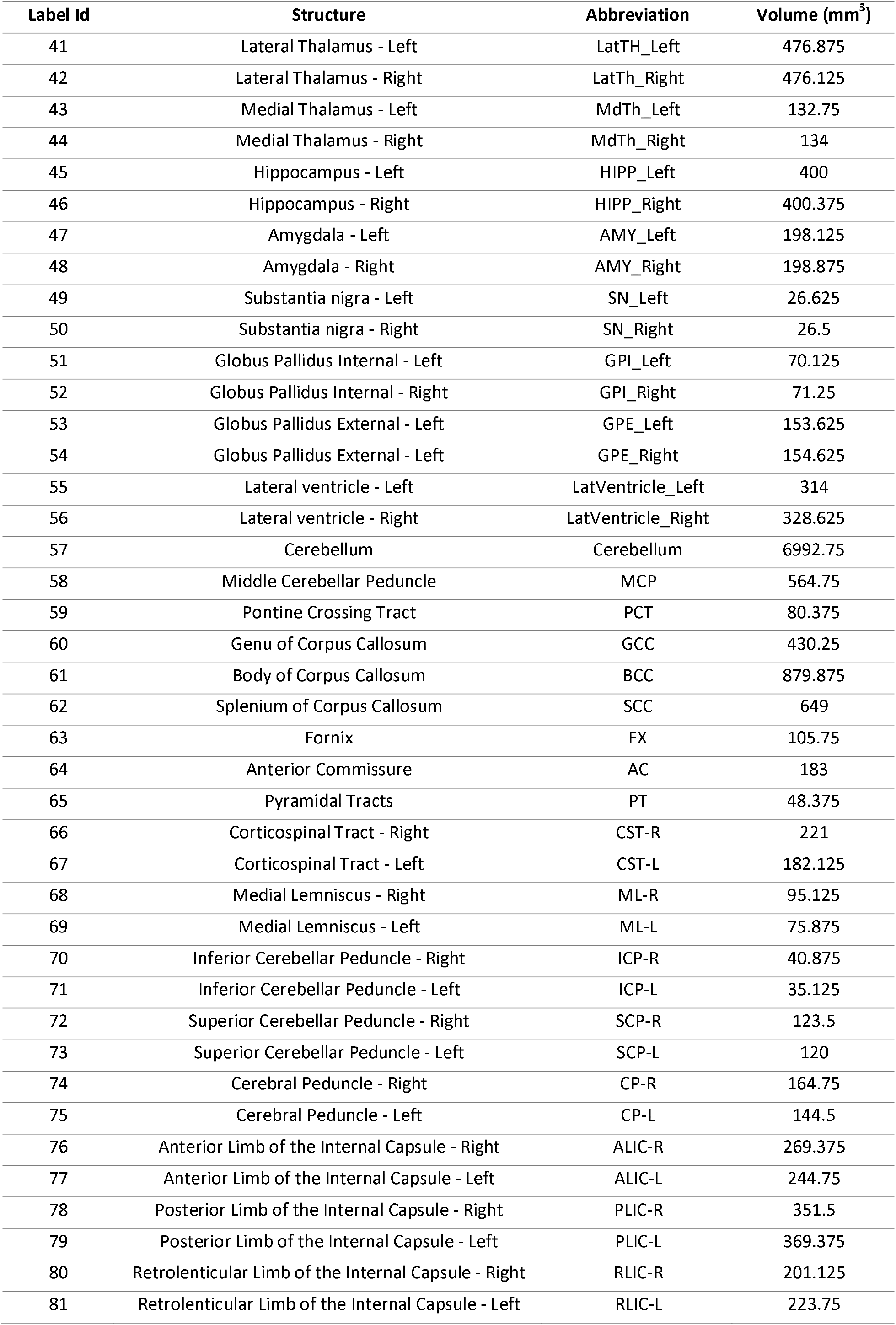
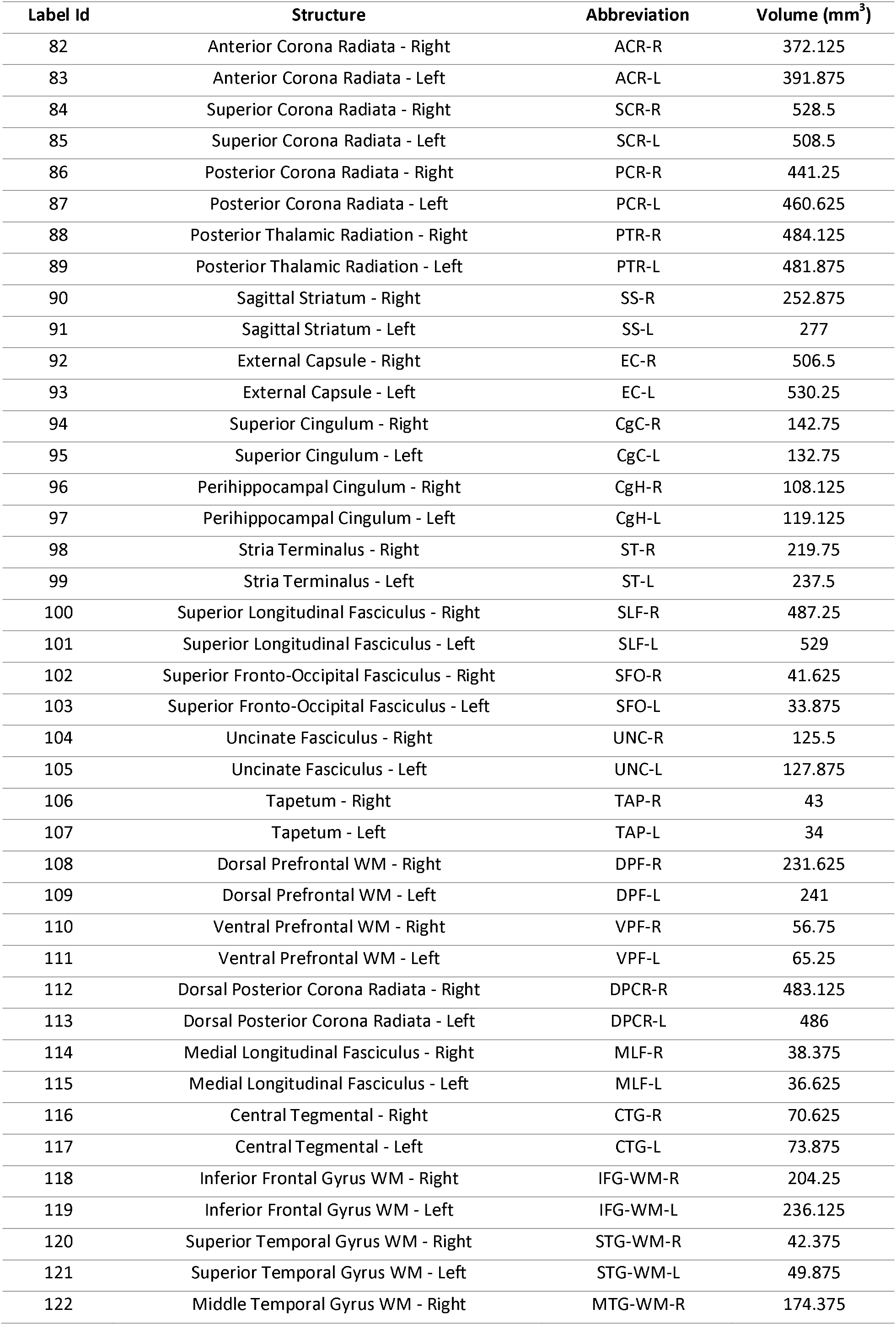
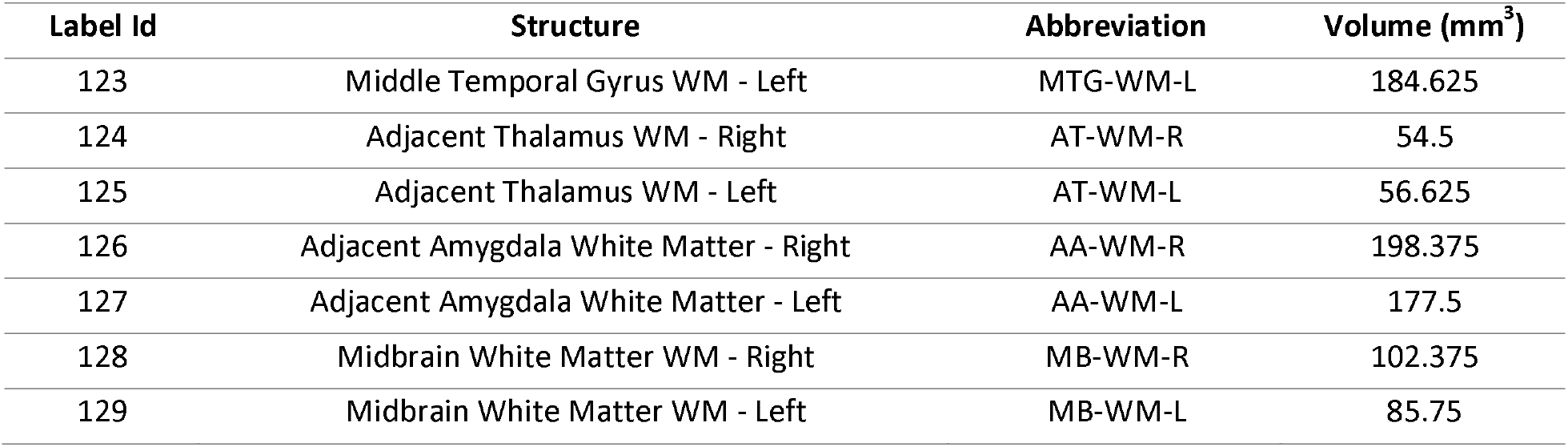
Description of ROIs in Combined GM/WM labelmap. We used ROIs defined by a published diffusion tensor-based rhesus macaque atlas to identify and segment WM tracts on our FA template (Zakszewski et al., 2014). These WM ROIs were combined with our GM ROIs (described in Table 2) into one consolidated labelmap that is described by this table, as well as the volume (in mm^3^) of each ROI in the labelmap.

To prevent motion artifacts and to ensure the safety of the animals, all monkeys were anesthetized for the duration of the scanning session. Anesthesia was initially induced with Ketamine HCI (15mg/kg IM) and the animals were subsequently intubated to allow for maintenance of anesthesia by inhalation of 1-2% isoflurane gas vaporized in 100% oxygen. Animals were then placed in a head first, supine position on the scanner bed, and their heads were immobilized in the head coil with foam padding. Heart rate and blood oxygenation levels were observed and recorded by trained veterinary staff throughout the duration of the procedure. Upon completion of the scans, the animals were extubated, returned to their housing environment, and their recovery monitored closely for several hours.

### 2.2 MRI acquisition procedures

Images were acquired with a Siemens Prisma whole body 3T MRI system (Erlangen, Germany) using a 16-channel pediatric head rf coil. A vitamin E tablet taped to the right side of the head served as a fiducial marker. The images acquired included 3D T1-weighted magnetization-prepared rapid gradient-echo (MP-RAGE) (Mugler and Brookeman, 1990), 3D T2-weighted sampling perfection with application optimized contrasts using different flip angle evolution (SPACE) (Mugler et al., 2000), and diffusion tensor imaging (DTI). Detailed parameters for each scanning sequence are described in the sections that follow.

#### 2.2.1 T1-weighted MP-RAGE

For 3D MP-RAGE imaging sequences, TE/TR/TI = 3.44/2600/913 ms, flip angle = 8°, voxel sizes were 0.5 mm isotropic and 224 slices were acquired. In-plane image sampling consisted of 320 by 320 data points in the phase-encoded and readout directions, respectively. Three MP-RAGE images were acquired in each imaging session and these were rigid-body registered to each other and averaged. Total MP-RAGE acquisition time was 31 minutes and 9 seconds.

#### 2.2.2 T2-weighted SPACE

For 3D SPACE imaging sequences, TE/TR = 385/3200 ms, flip angle = 120°, voxel sizes and the field of view were the same as the 3D MP-RAGE images described above. Three SPACE images were acquired in each imaging session and these were rigid-body registered to each other and averaged. Total SPACE acquisition time was 29 minutes and 42 seconds.

#### 2.2.3 Diffusion Tensor Imaging

Diffusion-weighted volumes were acquired using a monopoplar 2D diffusion-weighted, spin-echo echo planar imaging (EPI) sequence (TR/TE = 6700 ms/73 ms, GRAPPA factor = 2, echo train length = 52, resolution = 1 X 1X 1mm). Seven repetitions of 6 b0 volumes and 30 DW volumes with single b = 1000 s/mm^2^ were acquired with the phase-encoding direction being anterior to posterior (A -> P). To correct for the susceptibility induced distortions inherent to the EPI sequence, 1 b0 volume with opposite (P -> A) phase encoding directions was also acquired. Total diffusion MRI acquisition time was 29 minutes and 39 seconds.

### 2.3 MR image preprocessing

A set of preprocessing operations was first applied to all acquired MR images to correct various types of artifacts and to extract brain masks. For T1-weighted and T2-weighted anatomic images (MP-RAGE and SPACE images), the artifacts, such as motion and intensity inhomogeneity, were corrected first, and then the brains were extracted for template construction. For DTI, the distortion of phase-encoding, motion and eddy current artifacts were corrected first, and subsequently the templates were constructed.

#### 2.3.1 T1-weighted and T2-weighted anatomic image preprocessing

Both T1-weighted and T2-weighted images were preprocessed with identical procedures. In each imaging session, each of the three images were averaged after motion correction. For the motion correction, the first scanned image in each modality was selected as the reference, and other two images were registered to the reference using the rigid-body transformation. Next, the registered images and the reference image were averaged. All registration processing and averaging operations were carried out using ANTS (version 2.1; http://stnava.github.io/ANTs/) and FSL (version 5.0, http://fsl.fmrib.ox.ac.uk/fsl/fslwiki/). For all merged T1-weighed and T2-weighted images, intensity bias correction was performed using the “N4BiasFieldCorrection” tool in ANTS (Tustison et al., 2010). Next, T1- and T2-weighted head templates were constructed in the native coordinate using the “buildtemplateparallel.sh” tool in ANTS (Avants et al., 2010). Brain masks were drawn manually for head templates, and all bias field corrected subject T1- and T2-weighted images were nonlinearly registered to the corresponding head templates. With the resulting registration parameters, the template brain masks were inversely mapped to each subject T1- and T2-weighted image spaces and the brains were extracted by the brain mask. The process of intensity bias correction for all brain images was repeated using the “N4BiasFieldCorrection” tool, in which the subject brain mask was used to limit the correction region to improve correction quality.

#### 2.3.2 DTI preprocessing

All the DW volumes first underwent a de-noising step implemented in MATLAB (using a MATLAB script kindly provided by Dr. Sune Jespersen, Aarhus University) (Veraart et al., 2016). Next, a susceptibility-induced off-resonance field (h) was calculated from the 6 pairs of b0 volumes with opposite phase-encoding direction using “topup,” included in the FSL library (https://fsl.fmrib.ox.ac.uk/) (Andersson et al., 2003). An eddy current induced off-resonance field (e) and rigid-body transformations (r) between DW volumes to account for motion were estimated simultaneously using “eddy” (FSL). Finally, the 3 transformations (h, e, and r) were combined into one warp field to correct the de-noised DW volumes (Andersson and Sotiropoulos, 2016). “DTIFIT”, another tool included in FSL library, was used to fit the de-noised, corrected b0s and DW volumes to a single tensor (DTI) model.

### 2.4 Multi-modal template construction and atlas creation

After preprocessing operations for the anatomic and DTI data were completed, population-based average brain templates were constructed. The T2-weighted template was constructed first as the reference for the other template constructions as described below.

#### 2.4.1 T2-weighted template construction

All corrected T2-weighted brain images were initially aligned into a common space using a previously published T2 template (Calabrese et al., 2015) by rigid-body registration with the “antsRegistrationSyN.sh” tool. This reference frame was selected due to its high resolution and excellent tissue contrast, as well its orientation in the standard ‘MNI’ space (Frey et al., 2011). This way, all transformed brain images were represented in a commonly used reference frame, possessing the same orientation and position. Next, an ‘initial template’ was generated from the resulting images using the “buildtemplateparallel.sh” tool (https://github.com/ANTsX/ANTs/blob/master/Scripts/buildtemplateparallel.sh) with one iteration and affine registration, using the default similarity metric (probability mapping). Then, the newly generated ‘initial template’ was used as the target and the template building command was re-run in four iterations using fully deformable registrations. The output was the final T2-weighted template.

#### 2.4.2 T1-weighted template construction

The T2-weighted template was used as the reference for T1-weighted template construction. First, each corrected T1-weighted brain image was aligned to the corresponding T2-weighted brain image with a 12-parameter affine linear registration using mutual information as a cost function with the “antsRegistrantionSyN.sh” tool. Next, all T2-weighted brain images were b-spline non-linearly registered to the reference using the same software (ANTS version 2.1; http://stnava.github.io/ANTs/). Subsequently, with the resulting parameters, all aligned T1-weighted brain images were transformed to the T2-weighted brain template space. After averaging all transformed images, the T1-weighted brain template was obtained, which resided in the same reference frame as the T2-weighted template.

#### 2.4.4 DTI Template

The diffusion tensor template was constructed from all 18 individuals and resampled with 0.5 mm isotropic voxel spacing using DTI-TK (https://www.nitrc.org/projects/dtitk), a tensor-based spatial normalization tool (Zhang et al., 2007). The procedure described in Adluru et al. (2012) was followed, in which tensor-based registrations were performed to generate a diffusion MRI template, which was subsequently registered to the T2-weighted template. It has been demonstrated that templates created using tensor-based methods (i.e. DTI-TK) perform better than templates created using multimodal intrasubject registration methods (i.e. FA-T1 and B0-T2) (Adluru et al., 2012). For this reason, we adopted DTI-TK methods to generate the DTI template and then aligned the DTI template to the anatomical template space where multimodal registration operations were implemented only once, rather than to use intrasubject registrations to bring each subject’s DTI images into the space of the T2 before template construction. Once completed, the diffusion tensor template was non-linearly mapped to the T2-weighted template. To achieve this, the b0 template was first non-linearly registered to the T2w template using “antsRegistrationSyn.sh”, resulting in an affine transform and a warp field. Next, the diffusion tensor template was warped to the T2w space using the affine transform and the warp field generated from the previous step via “antsApplyTransforms” (with −e 2 option). Lastly, the affine transform and the warp field were combined into a single warp file which was used to reorient the diffusion tensors that were already in T2-weighted template space. The DTI-TK TVtool was then used to generate tensor derivatives (i.e. FA, MD, RD, and RGB color-coded FA images), which were subsequently reoriented with ITK-SNAP into RPI orientation to match the T1 and T2 templates.

### 2.5 Gray Matter(GM) Labelmap

The atlas of Saleem & Logothetis (2007) was used to define the boundaries of all cortical and subcortical Regions of Interest (ROIs) in the labelmap. ROIs were manually drawn on coronal sections of the T2 template by trained neuroanatomical experts (A.R.W., W.A.L., J.L.M.), using the 3D segmentation software ITK-SNAP (Yushkevich et al., 2006), and subsequently verified and edited in the axial and sagittal planes. **Table 2** describes the anatomical boundaries used to delineate this labelmap, and **Figure 1A** provides 3-dimensional renderings of the ROIs. Additionally, to facilitate neuroimaging modalities where scans have lower spatial resolution, we created an auxiliary version of the GM labelmap with condensed cortical and sub-cortical **ROIs** (**Table 2, Figure 1B.**)

**Figure 1:**
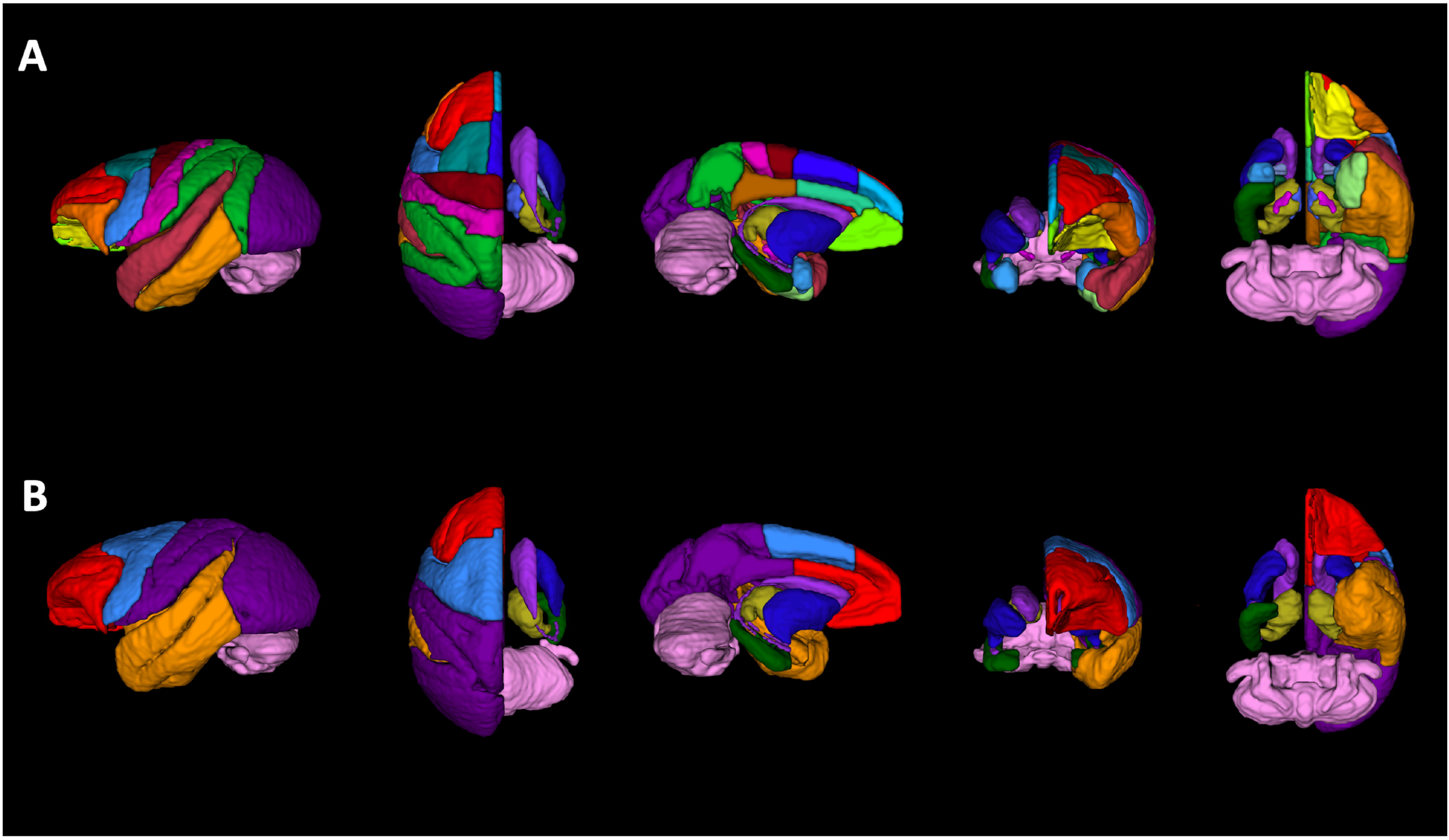
Rendering of labelmaps in 3-dimensions. (A) 3D-rendering of GM ROIs created using the 3D segmentation software ITK-SNAP, (Yushkevich et al., 2006). There are 57 cortical and subcortical ROIs defined in this labelmap. (B) Regions from the labelmap were subsequently categorized and combined to create a consolidated labelmap that defines 8 bilateral cortical ROIs and 9 subcortical structures, in order to facilitate neuroimaging modalities with low spatial resolution (i.e. rsfMRI, PET). To ease the visualization of subcortical structures, cortical regions in the right hemisphere were made transparent in this figure.

### 2.6 Combined GM/WM labelmap

To provide segmentations of white matter (WM), we used a published diffusion tensor-based rhesus macaque atlas (Zakszewski et al., 2014) to identify and segment WM tracts on our templates. In order to combine these WM ROIs with our gray-matter (GM) ROIs into one consolidated labelmap, the WM labelmap was first applied to the FA template. To accomplish this, the FA templaate from Zakszewski et al. (2014) was nonlinearly registered to our FA template, and the resulting transformation applied to the WM labelmap. This process resulted in a number of voxels being redundantly labeled as GM and WM structures. Using ITK-snap, the overlapping voxels were manually categorized by a trained observer (A.R.W.), and then labelmaps combined using FSL commands. The cerebellum was defined in this atlas using only one consolidated ROI and so all overlapping WM ROIs in this region were assigned to part of the cerebellar ROI.

## 3 RESULTS

We created a series of co-registered templates, including T1- and T2-weighted MRI as well as DTI, that are aligned with a set of labelmaps that define Regions of Interest (ROI) to facilitate the investigation of cortico- and thalamo-basal ganglia circuitry.

### 3.1 T1 and T2 templates and labelmaps

The anatomical T1 and T2 templates have identical native resolution (0.5 mm isotropic), and fields of view (60 mm by 79.5 mm by 50 mm). The templates are provided in neuroimaging informatics technology initiative (NIFTI) file format with all associated meta-data formatted to Brain Imaging Data Structure (BIDS) specifications (https://bids-specification.readthedocs.io/en/stable/), and have been submitted to the NITRC data exchange (www.nitrc.org) (5.9 MB). **Figure 2** illustrates axial and coronal views of the T1 **(Figure 2 A,C)** and T2 **(Figure 2 B,D)** templates, respectively, at multiple levels throughout the brain, with the labelmap for one hemisphere provided as an overlay. Our labelmap defines 57 ROIs (36 cortical regions and 21 subcortical structures) using the atlas of Saleem & Logothetis (2007) as a guide. All of the labels are bilateral, except for the cerebellum. For reference, the anatomical boundaries are described in **Table 2** and volumes of each ROI are reported in **Table 3.** Structures defined in this labelmap were chosen to facilitate the investigation of cortico-basal ganglia and thalamo-basal ganglia networks (Weiss et al., 2020a), and so less detailed segmentations are provided for regions such as the hindbrain and cerebellum. We also created a condensed version of the labelmap that is designed to be used for analyses where larger ROIs are useful, such as neuroimaging modalities with low spatial resolution like PET. This labelmap contains 17 ROIs (8 cortical regions and 9 subcortical structures), and is also described in **Table 2**. As in the more detailed labelmap, all labels in the condensed labelmap are bilateral except for the cerebellum.

**Figure 2:**
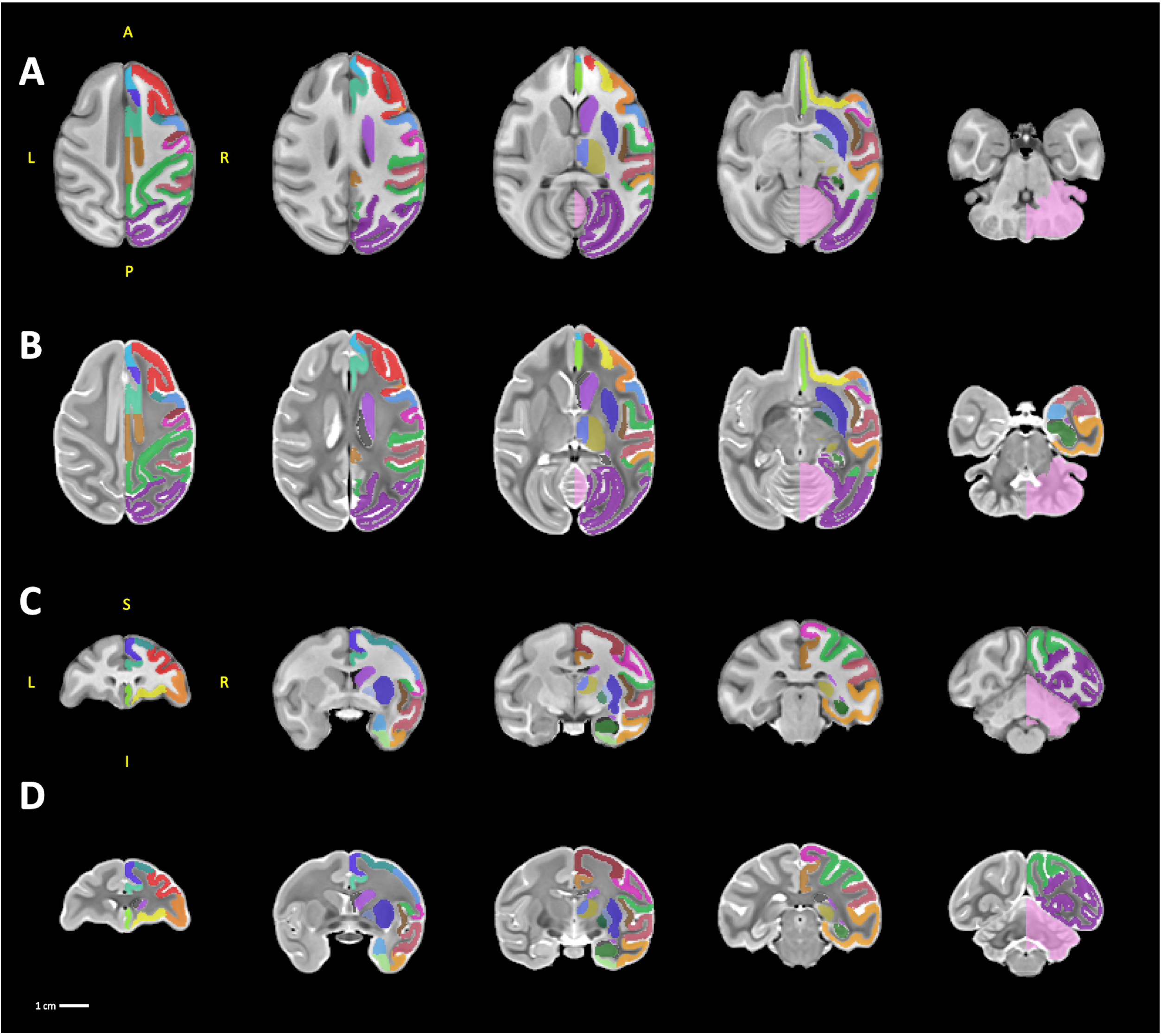
sMRI templates. A series of five axial and coronal sections displayed in neurological orientation through the T1 templates (A, C) and the T2 templates (B, D). For ease of viewing, the labelmap overlays are drawn only on the right hemisphere. *Abbreviations: A, anterior; I, inferior; L, left; P, posterior; R, right; S, superior.*

### 3.2 Diffusion templates/labelmaps

The diffusion tensor template was constructed independently from 18 individual DW volumes and registered to the shared T1 and T2 reference frame. The tensor template, its derivatives such as FA, MD, AD, and RD, were submitted in NIFTI file format, along with and BIDS formatted meta-data, for download on NITRC (www.nitrc.org) (74.7 **MB). Figure 3A, C** illustrates axial and coronal views of the FA template at multiple levels with the labelmap for one hemisphere provided as an overlay, and **Figure 3B, D** illustrates the RGB template as an overlay on the T1w template to highlight the spatial co-registration of the DTI and sMRI templates.

**Figure 3:**
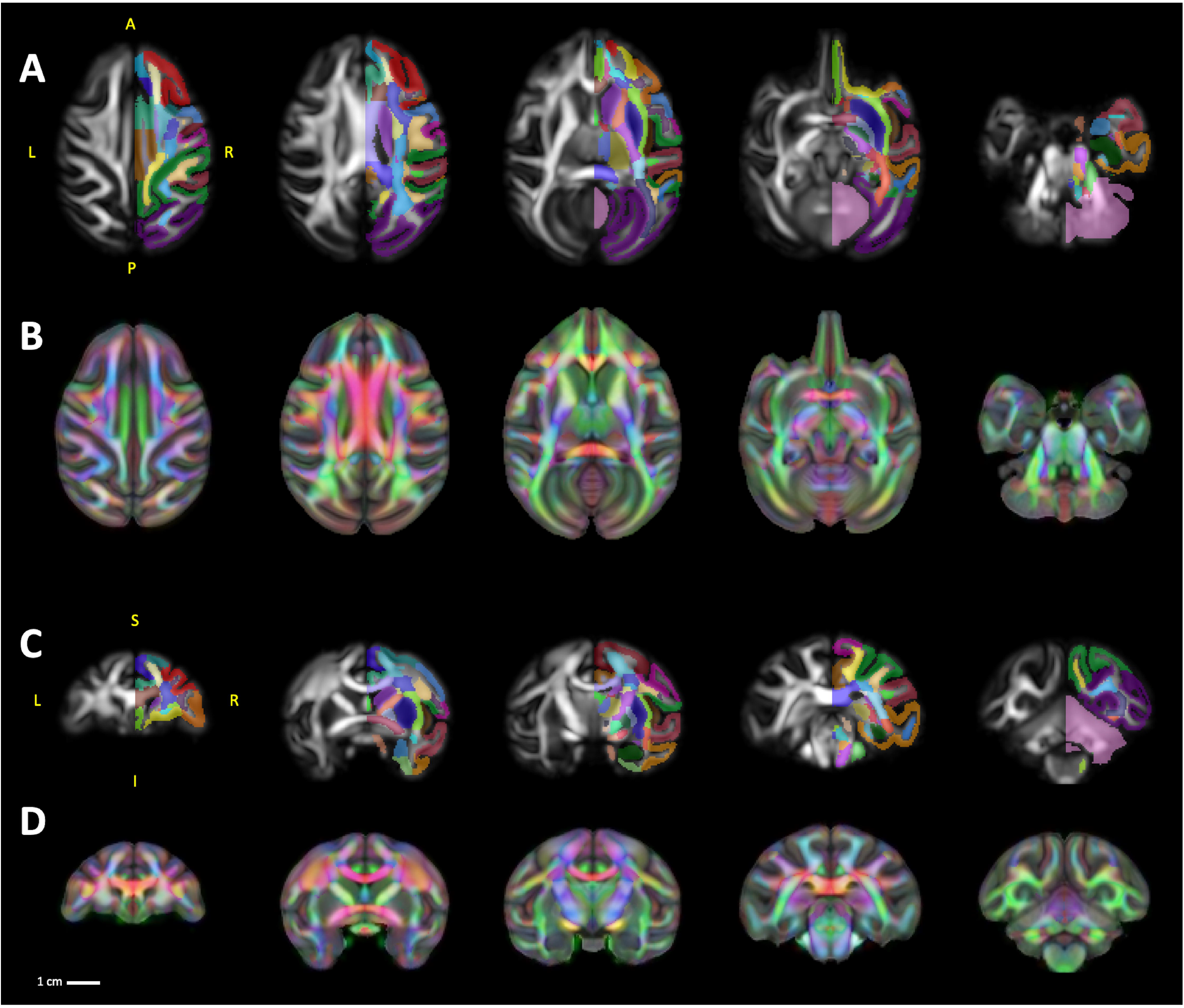
DTI Templates. A series of five axial and coronal sections displayed in neurological orientation through the FA template (A, C). For ease of viewing, the labelmap overlays are drawn only on the right hemisphere of the FA templates. To highlight the spatial co-registration of the DTI template with the sMRI templates (B, D) show visualization of RGB template as an overlay on the T1w template. *Abbreviations: A, anterior; I, inferior; L, left; P, posterior; R, right; S, superior.*

It is noteworthy that, because the template was built upon individual DW volumes with high resolution and good SNR, our FA template allows delineation of numerous cortical regions such as the dorsolateral prefrontal cortex and subcortical structures including those comprising the basal ganglia, in addition to white matter tracts throughout the brain. Therefore, to further broaden the applicability and take advantage of advancements in scanning technology, we created a labelmap that combines our GM segmentations **(Figure 4A)** with segmentations of 74 WM tracts defined by a previously published macaque DTI atlas (Zakszewski et al., 2014) in **Figure 4B**. After co-registration of both labelmaps to the template-space, we identified a small number of voxels that were labeled in both the GM and WM labelmaps **(Figure 4C, D**.) These voxels almost entirely fell along the boundaries between tissues (white matter-grey matter-CSF), **(Figure 4E, F**.) The resolution of our template, compared to the template by Zakszewski et al. (2014), as well as differences in the age ranges of the animals in each template, required us to refine the boundaries of these ROIs to better match the anatomy. **Table 3** describes all of the ROIs in this labelmap, and reports the volumes of each.

**Figure 4:**
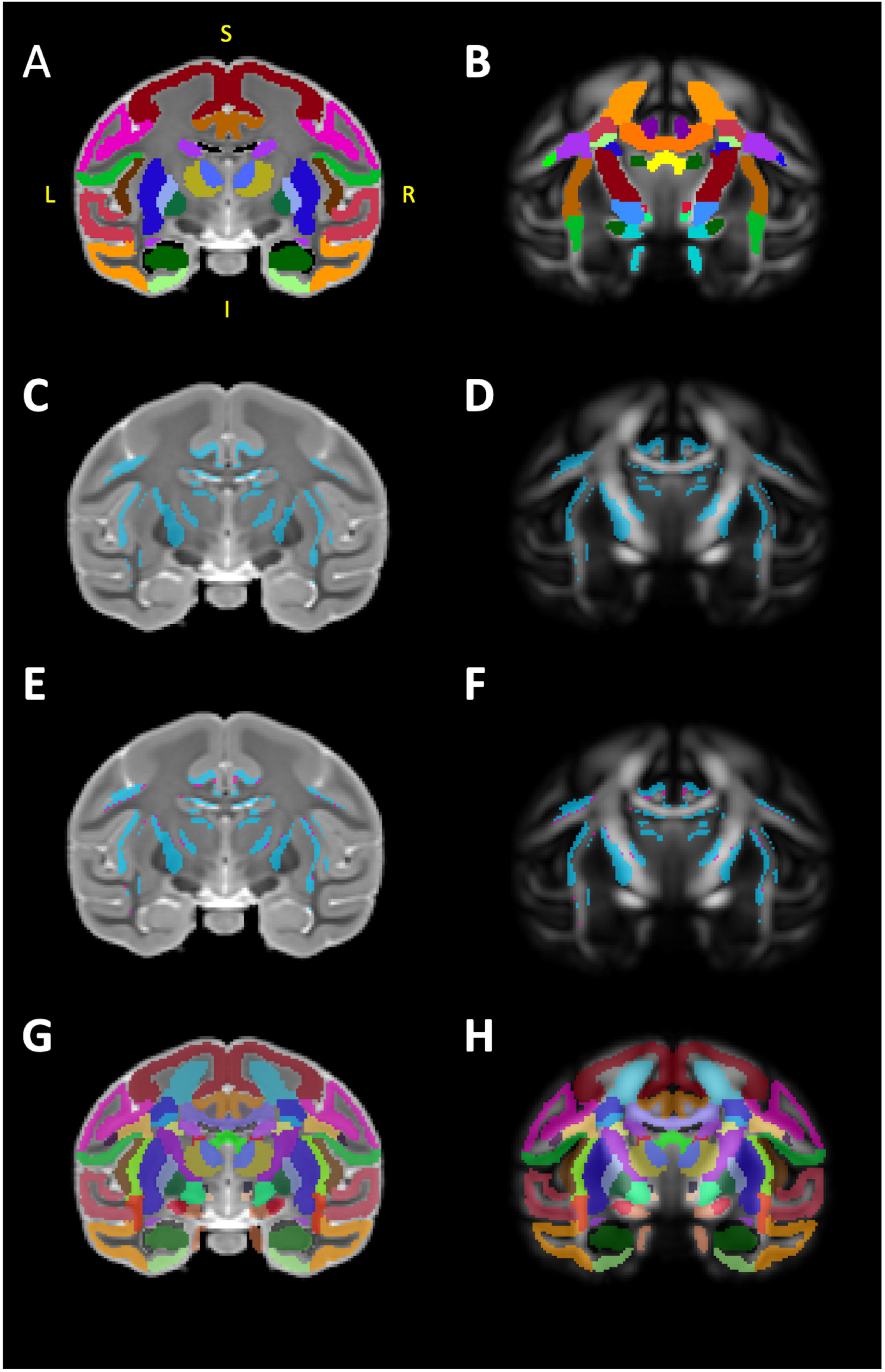
Combing gray and white matter labelmaps. (A) illustrates our gray matter (GM) labelmap overlaying the T2 template. We used a published diffusion tensor-based rhesus macaque atlas (Zakszewski et al., 2014) to identify and segment white matter tracts on our FA template, illustrated in (B). In order to combine these two labelmaps, using FSL tools, we first identified overlaps (i.e. double-labeled voxels) between the WM labelmap and our GM labelmap, these voxels are illustrated in blue in (C) and (D). ITK-SNAP was used to manually categorize these voxels as belonging to WM or GM, and the results of this classification is illustrated in (E) and (F), with blue voxels categorized as GM and magenta voxels categorized as WM. Finally, the categorized voxels were re-assigned to the appropriate GM/WM labelmap and ROI, and the two labelmaps were then consolidated into one. The results of this combination are illustrated in (G) and (H), overlaying the T2 and FA templates. *Abbreviations: I, inferior; L, left; P, posterior; R, right.*

### 3.2 Example Applications

We have included several examples of scans collected from individual macaques using a variety of neuroimaging modalities (sMRI, DTI, PET) with co-registered labelmaps (generated by AntsRegistration) to demonstrate potential applications for the ONPRC18 atlas. First, as part of an ongoing longitudinal study of cortico-basal ganglia circuitry in a rhesus macaque model of Huntington’s disease, the atlas was applied to a DTI scan acquired from one of the individuals used to build the template after delivery of the mutant *HTT* gene to the caudate and putamen **(Figure 5A**.) Close manual inspection of the overlays revealed satisfactory results of the registration and accurate anatomical segmentations. To further probe the versatility of this atlas, we also applied it to structural scans (T1w/T2w) that were collected from a juvenile, 3-year old Japanese macaque (Macaca fuscata) bearing a CLN7 gene mutation (macaque model of Batten disease) **(Figure 5B,C)** again with satisfactory registration despite differences in macaque species, age and disease model. Finally, we have applied our consolidated labelmap to 18F-Fallypride PET and 18F-FDG PET images **(Figure 5 D,E**, respectively) collected from naïve adult rhesus macaques with satisfactory registration, despite the lower spatial resolution of these scans. Both PET scans were collected with co-aligned CT, and the atlas was registered using the CT skull as a reference and then moved into PET space. These examples demonstrate some of the potential applications for this atlas, and suggest that it can be a reliable and versatile tool for different kinds of macaque neuroimaging studies.

**Figure 5:**
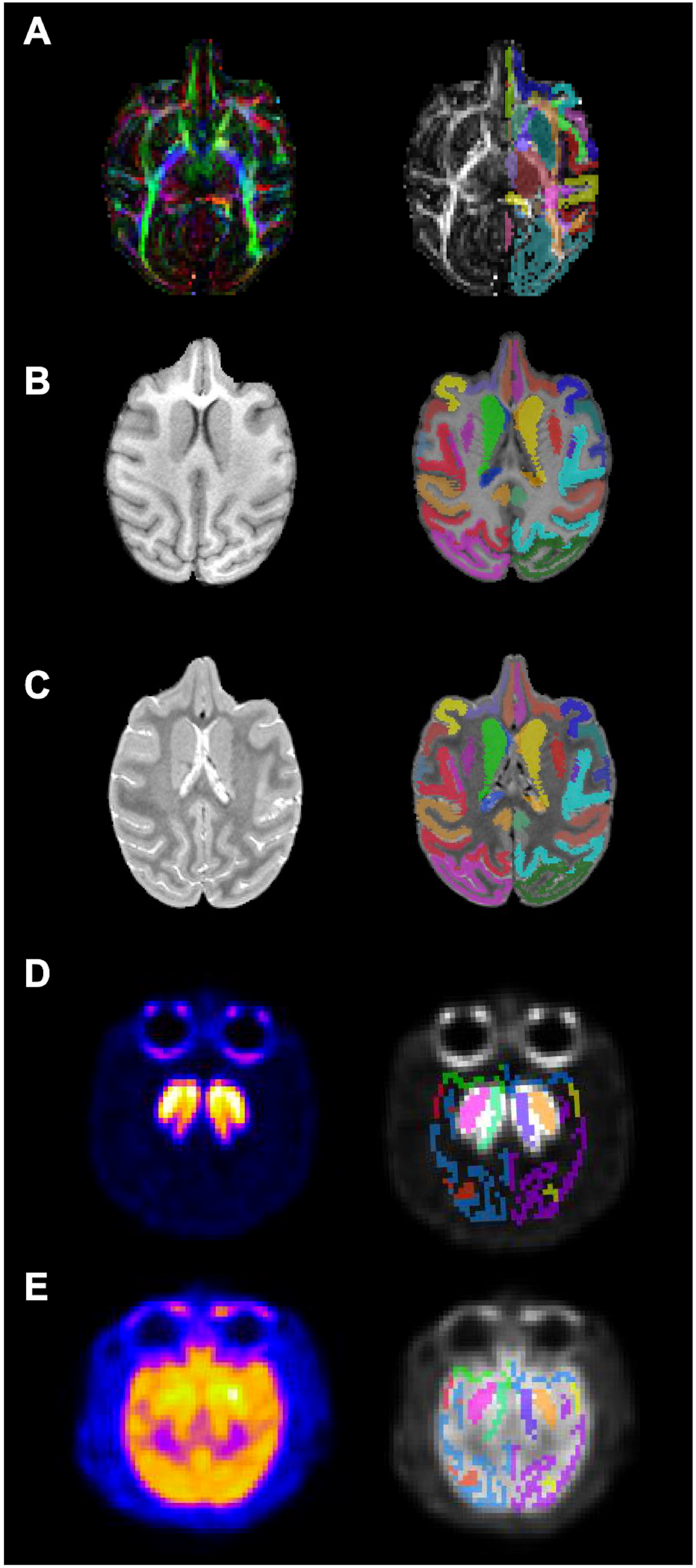
Example application of multimodal templates. A series of axial sections with co-registered labelmaps that illustrate applications of the ONPRC18 atlas to scans collected from individual macaques using a variety of neuroimaging contrasts and modalities (DTI, sMRI, PET). Label maps were generated by AntsRegistration software version 2.1; (http://stnava.github.io/ANTs/). (A) DTI scan acquired from one of the individuals in the template at a later timepoint. (B, C) sMRI scans (T1w/T2w) that were collected from a juvenile (3yo) Japanese macaque (Macaca fuscata). (D, E) 18F-Fallypride PET and 18F-FDG PET in a naïve female rhesus macaque. These examples demonstrate some of the potential applications for this atlas, and suggest that it can be a reliable and versatile tool for a wide variety of different kinds of macaque neuroimaging studies.

## 4 Discussion

We created a brain atlas designed to facilitate studies with macaques employing both structural MRI and diffusion weighed imaging (DWI). Template images of the structural contrasts (T2/SPACE, T1/MP-RAGE) and DWI parameter maps (FA, AD, MD, RD, RGB, b0) were constructed in the 0.5mm isotropic voxel-spacing reference frame. The labelmap provides segmentations of 57 gray matter ROIs (36 cortical regions and 21 subcortical structures) as well as 74 white matter tracts, and overlays all of the structural and diffusion templates. A condensed version of the labelmap was also created, with 17 larger ROIs, to facilitate imaging modalities with lower spatial resolution, such as fMRI and PET.

The templates here were created from in vivo scans, rather than scans of post-mortem fixed-tissue as in Calabrese et al. (2015); Feng et al. (2017); Reveley et al. (2017). The obvious merit to the latter is the ability to employ very lengthy scanning sequences (that would be inappropriate for use with living animals) in order to collect high quality data. Yet, the scanning acquisition parameters employed here resulted in templates with image quality approaching that of many post-mortem datasets. Furthermore, fixation techniques used on post-mortem tissue can alter brain volume and water content, or induce non-rigid distortions, and therefore represent an important limitation in the application of post-mortem atlases to *in vivo* neuroimaging, and to DWI in particular. In contrast, the templates included here provide references that were acquired from living subjects, a feature that was shown to result in good registration accuracy when applying this atlas to other *in vivo* neuroimaging studies with macaques.

The construction of the ONPRC18 templates contrasts from previously available NHP atlases in several additional significant ways. First, rather than mirroring across the midline to create a laterally symmetric template as in Calabrese et al. (2015); Moirano et al. (2019), the ONPRC18 atlas preserved bilateral asymmetry between hemispheres in order to better reflect variability among the macaque population, as suggested by Frey et al. (2011). Second, the demographic distribution of the population ONPRC18 animals is in the middle-age range for macaques (8.1 ± 2.2 years) and includes a higher proportion of females (66%) than many existent population-based templates for adult monkeys: *INIA19 template:* 8 ± 1.6 years, all males (Rohlfing et al., 2012); *NMT template:* 5.5y ± 1.7 years, 19% female (Seidlitz et al., 2018); *MNI template:* ‘young adult’, 20% female (Frey et al., 2011); *112RM-SL template:* 19.9 ± 6.9 years, 27% females (McLaren et al., 2009); versus *UNC-Wisconsin Neurodevelopment template: covers 0-3 years, 50% females (Young et al., 2017); UNC-Emory Infant template:* covers 0-12months, 50% females (Shi et al., 2016). These differences may be particularly relevant for brain regions and structures with a protracted development that extends beyond puberty or with vulnerability to advancing age, as well as for cross-sectional studies investigating sex differences. Third, new advances in SPACE pulse sequences have made it possible to collect high quality T2w images at identical resolution as T1w MPRAGE images, enabling either contrast to be used as a base anatomical dataset. Given that the T2w contrast provides a better reference for registration of DWI images than T1w (Adluru et al., 2012), the T2w template was selected as the primary reference for the construction of the ONPRC18 DTI- and T1w-templates. Finally, we noted that the contrast of the SPACE images offered some advantages for the anatomical delineation of deep-brain structures surrounded by white matter, such as the basal ganglia and thalamus, as compared to MPRAGE (see **Figure 2**). Given the relevance of these structures to the circuitry defined in the labelmap, we assigned the T2w template as the primary reference.

The ONPRC18 labelmap includes gray matter ROIs that were hand drawn with reference to the cortical and subcortical boundaries described by Saleem and Logothetis (2007). Building on previous work from our group, the INIA19 atlas (Rohlfing et al., 2012), we have made several practical improvements in the labelmap to facilitate in vivo multi-modal neuroimaging studies. First, by manually defining every GM ROI on a template generated from 18 animals, rather than transferring boundaries from a series of 2D histological sections defined on a single animal, we avoided the limitations inherent to generating 3D digital atlases from 2D histological drawings, as described by Reveley et al. (2017); Moirano et al. (2019). This resulted in a labelmap that is well matched to the image resolution of in vivo MRI and quite tolerant of individual variation in neuroanatomy, a feature clearly demonstrated in **Figure 5**. Similarly, by manually drawing ROIs on both hemispheres of the template rather than mirroring the labelmap across the midline, the ability to describe individual variabilities in hemispheric asymmetry is improved. Lastly, the ONPRC18 labelmap defines many subcortical structures such as the caudate, putamen, internal and external globus pallidus, substantia nigra, as well as lateral and medial subdivisions of the thalamus. This feature contrasts with the widely used D99 labelmap that combines the caudate, putamen, and nucleus accumbens into one striatal ROI, (Reveley et al., 2017; Seidlitz et al., 2018). For these reasons, the ONPRC18 atlas will fill an unmet need for NHP imaging resources suitable for studies of cortico-basal ganglia and thalamo-basal ganglia circuitry.

To further increase the relevance of this tool for investigations of cortico- and thalamo-basal ganglia, we enhanced the ONPRC18 GM labelmap by combining it with segmentations of WM regions defined in a previously published macaque DTI atlas (Zakszewski et al., 2014). The combined GM-WM labelmap eliminates the need for researchers to apply separate GM and WM labelmaps to their datasets, thereby reducing the number of transformations needed since only one registration will be required. Additionally, the increased precision created by the elimination of overlaps in the labels, see **Figure 4**, enables the same gray and white matter regions to be more consistently identified. This is a unique feature of this atlas, and it will improve the validity of comparisons between scans acquired from the same subject in different imaging modalities.

We are currently applying the ONPRC18 atlas to a longitudinal study querying cortico- and thalamo-basal ganglia circuitry in a newly developed AAV-mediated rhesus macaque model of Huntington’s disease (HD) (Weiss et al., 2020a). In this context, we have found preliminary evidence of DTI changes occurring early in disease progression that correlate with cognitive and motor disease phenotypes (Weiss et al., 2020b), progressive reduction in putamen volume (unpublished data), and have begun to quantify changes in regional dopamine receptor binding potentials with F18-Fallypride PET (unpublished data, **Figure 5**.) Additional work is also underway using this atlas to assess regional brain atrophy and alterations in white matter microstructure in a Japanese macaque model of Batten’s disease (McBride et al., 2018), demonstrating that these templates have wide versatility and can be successfully applied to other species and ages of macaques.

In summary, it is our hope that this new ONPRC18 atlas with corresponding label maps will serve as an updated resource for NHP researchers to facilitate investigations of macaque brain circuitry, for developing and characterizing models of neurological disease, assessing the pre-clinical safety and biodistribution of therapeutics and/or developing imaging signatures of disease as outcome measures for pre-clinical trials.

## ACKNOWLEDGEMENTS

We sincerely thank the entire ONPRC Division of Comparative Medicine for the outstanding care of our rhesus macaques, with special acknowledgement of the efforts that Lauren Drew Martin, Theodore Hobbs, Michael Reusz, Brandy Dozier, Alona Kvitky, and Isabel Bernstein contributed to this work. This research was supported by NIH/NINDS R01NS099136, NIH/NINDS R24 NS104161 (NHP Resource for Preclinical Studies of Neurodegenerative Disease), NIH/NINDS F32NS110149-01, NIH/NIAAA U24AA025473, the Bev Hartig Huntington’s Disease Foundation and a grant from the M.J. Murdock Charitable Trust.

